# Effects of low temperature and cold-acclimation on photoinhibition and singlet oxygen production in four natural accessions of *Arabidopsis*

**DOI:** 10.1101/777060

**Authors:** Heta Mattila, Kumud B. Mishra, Iiris Kuusisto, Anamika Mishra, Kateřina Novotná, David Šebela, Esa Tyystjärvi

## Abstract

To understand the effects of low temperature and cold-acclimation on reactive oxygen species and photoinhibition of photosystem II (PSII), light-induced inactivation of PSII was measured at 22 and 4 °C from four *Arabidopsis thaliana* accessions (Rschew, Tenela, Columbia-0 and Coimbra) grown under optimal conditions. Photoinhibition was also measured at 4 °C from plants cold-acclimated at 4 °C for two weeks. Measurements were done in the absence and presence of lincomycin that blocks PSII repair, and PSII activity was assayed with the ratio of variable to maximum chlorophyll *a* fluorescence (F_V_/F_M_) and with light-saturated rate of oxygen evolution using a quinone acceptor. Of the non-acclimated accessions, Rschew was the most tolerant to photoinhibition and Coimbra the least; the rate constants of photoinhibition of the most sensitive accession were 1.3-1.9 times as high as those of the tolerant ones. The damaging reaction of photoinhibition in non-acclimated plants was slower or equal at 4 °C than at 22 °C. The rate constants of photoinhibition of cold-acclimated plants, at 4 °C, were 0.55 to 1.25 times as high as those of non-acclimated plants; the protective effect of cold-acclimation on photoinhibition was consistent in Columbia-0 and Coimbra whereas Rschew and Tenela were either slightly more tolerant or susceptible, depending on the method used to assay photoinhibition. Production of singlet oxygen, measured from thylakoid membranes isolated from non-acclimated and cold-acclimated plants, did not decrease due to cold-acclimation, nor did singlet oxygen production correlate with the rate of photoinhibition or with flavonol contents of the leaves.

## 1. Introduction

Light is the energy source of plants, but Photosystem II (PSII) is also damaged by light, and synthesis of a new D1-protein is needed for the recovery of the electron transfer activity of PSII (for reviews, see Tyystjärvi 2013; Nath et al. 2013). The initial rate of the damage is directly proportional to the intensity of light (Tyystjärvi and Aro 1996), and when damage is faster than repair, e.g. under high light, non-functional PSII units accumulate. In the literature, the term “photoinhibition” has been used to describe several phenomena; here we strictly refer to the light-induced irreversible loss of activity of PSII. The molecular mechanism of the damage to PSII is still under debate (Tyystjärvi 2013).

Low temperature is one of the major factors limiting growth and geographical distribution of plant species. The repair of PSII slows down if temperature drops (Greer et al. 1986; Salonen et al. 1998; Grennan and Ort 2007). Results about the effect of temperature on the rate of the damage itself, however, vary. A decrease in the rate of damage with a decrease in temperature in pumpkin thylakoids (Tyystjärvi et al. 1994; Tyystjärvi 2013) and chloramphenicol-treated leaves (Tyystjärvi 1993), lack of a clear temperature dependence in the cyanobacterium *Synechocystis* (Allakhverdiev and Murata 2004), and an increase in the rate of damage with decreasing temperature in the leaves of cucumber (Sonoike et al. 1999), *Chenopodium album* (Tsonev and Hikosaka 2003) and cotton (Kornyeyev et al. 2003) have been reported. According to the “excitation pressure” hypothesis (Sonoike et al. 1999; Kornyeyev et al., 2003) photoinhibition is expected to increase with decreasing temperature because light absorption and subsequent charge separation are almost temperature-independent but the sink capacity decreases with temperature because carbon fixation slows down (Abat and Deswal 2009; for a review, see Yamori et al. 2014). The “over-reduction” of electron transfer chain may increase singlet oxygen (^1^O_2_) production because reduced electron acceptors promote PSII charge recombination reactions. ^1^O_2_, in turn, could then damage PSII (see e.g. Triantaphylidès and Havaux 2009; Vass and Cser 2009). Photosystem I (PSI) is usually less susceptible to light-induced damage than PSII (Tyystjärvi et al. 1989), but also photoinhibition of PSI was reported in some cold-susceptible species at low temperatures (Sonoike et al. 1999).

Prolonged exposure to low but non-freezing temperatures triggers cold-acclimation in several plant species, causing large changes in gene expression and modifying multiple physiological processes, including synthesis of protective substances, changes in the composition of membrane lipids (e.g. Murata et al. 1992), and changes in enzyme activities or amounts of other substances protecting against reactive oxygen species (ROS). Eventually cold-acclimation allows them to survive or grow at even lower temperatures than required for acclimation, but the improvement of the cold-tolerance is species-specific (for a review, see e.g. Theocharis et al. 2012; Crosatti et al. 2013). Light plays an important role in the development of full cold-acclimation (Soitamo et al. 2008), and it has been suggested that plants respond to coldness partly through sensing and responding to the reduced electron transfer chain (Gray et al. 1996; Ivanov et al. 2006; Crosatti et al. 2013).

Cold-acclimation (Somersalo and Krause 1989; Gray et al. 1996; Krause et al. 1999; Savitch et al. 2000; Venema et al. 2000; Sane et al. 2003; Shang et al. 2003) or over-expression of cold-inducible genes (Yang et al. 2010) have been reported to attenuate photoinhibition of PSII at low temperatures. In some species this is due to increased activity of the repair cycle of PSII (Krause et al. 1999; Venema et al. 2000; Shang et al. 2003; Grennan and Ort 2007; Rogalski et al. 2008) but also the rate of the damaging reaction of photoinhibition has been reported to diminish due to cold-acclimation (Vonshak and Novoplansky 2008). The protection has been hypothesized to be based on the ability of cold-acclimated plants to keep the Q_A_ electron acceptor of PSII more oxidized in the light even at low temperature (Öquist et al. 1993; Gray et al. 1996). Cold-acclimation can increase the activities of the enzymes of the Calvin-Benson cycle, which leads to an increase in the rates of carbon fixation at low temperatures (Hurry et al. 1994; Strand et al. 1999). Alternative electron transfer routes (cyclic electron transfer and electron transfer to the plastid terminal oxidase) may also function more efficiently after cold-acclimation (e.g. Ivanov et al. 2012; Mishra et al. 2019). In many plant species, cold-acclimation also leads to changes in the redox potentials of the electron transport chain of PSII, possibly modifying recombination reactions and ^1^O_2_ yield (Janda et al. 2000; Ivanov et al. 2001; Sane et al. 2003).

The amounts of xanthophyll pigments and/or non-photochemical quenching of excitation energy (NPQ) can increase during cold-acclimation (e.g. Krause et al. 1999; Venema et al. 2000). In the case of cold-tolerant plants, protection by NPQ against low-temperature-induced photoinhibition may, however, be important only in short-term cold-stress (Havaux and Kloppstech 2001). Furthermore, cold-acclimation may affect the concentrations of anthocyanins and flavonols. Although in the vacuole, flavonols have been found in chloroplasts of several species (Saunders and McClure 1976). In *Phillyrea latifolia* chloroplast-envelope-located flavonols were reported to quench ^1^O_2_ (Agati et al. 2007). Flavonols are preferentially located at the lipid-water interphase (Scheidt et al. 2004), which allows them to quench ^1^O_2_ produced within the membrane. These properties may make flavonols important scavengers of ^1^O_2_, as the lifetime of ^1^O_2_ in plant cells is so short (for reviews, see Mattila et al. 2015; Arellano and Naqvi 2016) that the damage caused by ^1^O_2_ is expected to occur near the site of origin of this ROS (Moan 1990).

*Arabidopsis thaliana* grows over a broad geographic range with varying temperatures, and therefore, effects of low temperature and cold-acclimation on photoinhibition can be investigated in natural accessions of this model plant species. In the present study, we used four accessions (Rschew, Tenela, Columbia-0 and Coimbra) with different cold-acclimation capacities. Previous investigations on photoinhibition of PSII at low temperatures have mostly been conducted by illuminating plants in the absence of a translation inhibitor (Gray et al. 1996; Krause et al. 1999; Venema et al. 2000; Sane et al. 2003). Thus, it is not clear whether the observed changes in photoinhibition depend on differences in the rate of damage or repair. Moreover, previous reports investigating the effect of cold-acclimation on photoinhibition of PSII have mostly used only chlorophyll *a* fluorescence to assay photoinhibition (e.g. Gray et al. 1996; Krause et al. 1999; Venema et al. 2000; Sane et al. 2003). In the present study, to differentiate between NPQ, repair and damage, we illuminated leaves in the presence and absence of the chloroplast translation inhibitor, lincomycin, and assayed photoinhibition by chlorophyll *a* fluorescence as well as by oxygen evolution. In addition, ^1^O_2_ production was measured to understand the effects of cold-acclimation and this ROS on photoinhibition.

## 2. Materials and methods

### 2.1. Growth conditions

*A. thaliana* accessions, Rschew, Tenela, Columbia-0, and Coimbra, were grown in a growth chamber (FytoScope FS-RI 1600, Photon Systems Instruments, Brno, Czech Republic) for six weeks at day/night temperatures of 21 °C/18 °C, with day/night light rhythm of 8 h/16 h (photosynthetic photon flux density, PPFD, of 100 μmol m^−2^ s^−1^) with ~60 % humidity, as described in Mishra et al. (2014). After six weeks of growth, half of the plants were shifted for cold-acclimation at 4 °C for two weeks (ACC) while the rest of the plants (NAC) were kept in the same conditions as before.

### 2.2. Pigments

Chlorophyll and flavonol contents were measured from intact leaves with a nondestructive handheld device (Dualex Scientific, Force-A, Paris, France) after seven to eight weeks of growth. At least three individual uniform-sized NAC and ACC plants of each accession were selected, from which three fully expanded leaves were measured.

### 2.3. Gas exchange measurements

Net CO_2_ assimilation rates of individual attached leaves of NAC and ACC plants of each accession, after seven to eight weeks of growth, were measured with a gas exchange measuring system LI-6400-17 (Li-Cor, Biosciences, Linkoln, Nebraska, USA) using a 6400-15 *Arabidopsis* chamber of aperture diameter of 1 cm (Li-Cor, Biosciences, Linkoln, Nebraska, USA). CO_2_ concentration in the chamber was set to 385 ppm, air humidity to 60 ± 5 % and temperature to 22 °C. Light-acclimated leaves were illuminated first for 2 min at the PPFD of 100 μmol m^−2^ s^−1^ and then for 45 min at the PPFD of 2000 μmol m^−2^ s^−1^.

### 2.4. Photoinhibition measurements

After eight weeks of growth, detached leaves, with petioles under water, were illuminated for 0–45 min at the PPFD of 2000 μmol m^−2^ s^−1^ at 22 °C or 4 °C in a growth chamber (FytoScope FS 130, Photon Systems Instruments, Brno, Czech Republic). To prevent the repair cycle of PSII, leaf petioles were incubated overnight in lincomycin (0.4 mg/ml) solution, under the low irradiance of ~10 μmol m^−2^ s^−1^. Before and after the treatment, leaves were kept for 30 min in dark at 22 °C, after which the chlorophyll *a* fluorescence parameter, F_V_/F_M_ (variable to maximum fluorescence), was measured by Handy Fluorcam FC 1000-H (Photon System Instruments, Brno, Czech Republic). After the fluorescence measurements, thylakoid membranes from 3–6 leaves were isolated as described by Hakala et al. (2005) and immediately stored at −80 °C. PSII activity was then measured with a method parallel to fluorescence, by measuring the light-saturated rate of oxygen evolution from the thylakoid membranes with an oxygen electrode (Hansatech, King’s Lynn, UK) as described by Hakala et al. (2005) using 0.5 mM 2,6-dimethylbenzoquinone as an electron acceptor. Chlorophyll concentration of isolated thylakoid membranes was measured according to Porra et al. (1989). Whenever lincomycin was used, the rate constant of photoinhibition (k_PI_) was calculated by fitting the decrease in F_V_/F_M_ or in the rate of oxygen evolution, as indicated, to the first order reaction equation.

### 2.5. ^1^O_2_ and thermoluminescence measurements

The rate of ^1^O_2_ production of isolated thylakoid membranes (100 μg chl/ml) in high light (PPFD 4000 μmol m^−2^ s^−1^), at 4 °C or at 22 °C, was estimated by measuring the consumption of oxygen by 20 mM histidine (Telfer et al. 1994, Rehman et al. 2013) as described in Karonen et al. (2014).

Thermoluminescence bands were recorded with a luminometer from isolated thylakoid membranes (500 μg chl/ml) as described earlier (Tyystjärvi et al. 2009) in the presence or absence of 20 μM 3-(3,4-dichlorophenyl)-1,1-dimethylurea (DCMU), as indicated. The thylakoid membranes for ^1^O_2_ and thermoluminescence measurements were isolated as described above from NAC and ACC plants taken directly from growth chambers.

### 2.6. Statistical tests and figures

Significances of differences were tested by calculating Student’s t-test (two-tailed, unequal variances). Fitting of photoinhibition to the first order reaction equation were done in SigmaPlot (Systat Software, Inc) as well as all figures.

## 3. Results

### 3.1. Assaying photoinhibition of PSII, in the presence of repair, by chlorophyll a fluorescence

Detached leaves from non-acclimated (NAC) *A. thaliana* accessions (Rschew, Tenela, Columbia-0 and Coimbra) were given a high light treatment (PPFD of 2000 μmol m^−2^s^−1^) either at their growth temperature (at 22 °C) or at 4 °C. We measured maximum quantum yield of PSII photochemistry (F_V_/F_M_; after 20 min of dark-incubation) during the 0–45 min high light treatment (Fig. 1A–B). Illumination was conducted in the absence of lincomycin to allow the repair of the D1-protein to proceed simultaneously with photoinhibitory damage to PSII. In addition, leaves from plants cold-acclimated (ACC) for two weeks at 4 °C were illuminated under the same high light at 4 °C (Fig. 1C). Of the four accessions, Coimbra and Rschew seemed the most susceptible and tolerant to photoinhibition of PSII, respectively (Fig. 1). In NAC plants, on the average, F_V_/F_M_ values declined faster at 4 °C than at 22 °C (Figs. 1A–B), although the difference was statistically significant only for Tenela after 15 min (*P* = 0.03) and 30 min (*P* = 0.01) and for Rschew after 45 min (*P* = 0.01) of the light treatment. However, the rate of photoinhibition at 4 °C did not differ significantly between NAC and ACC plants in any of the accessions.

**Fig. 1.**
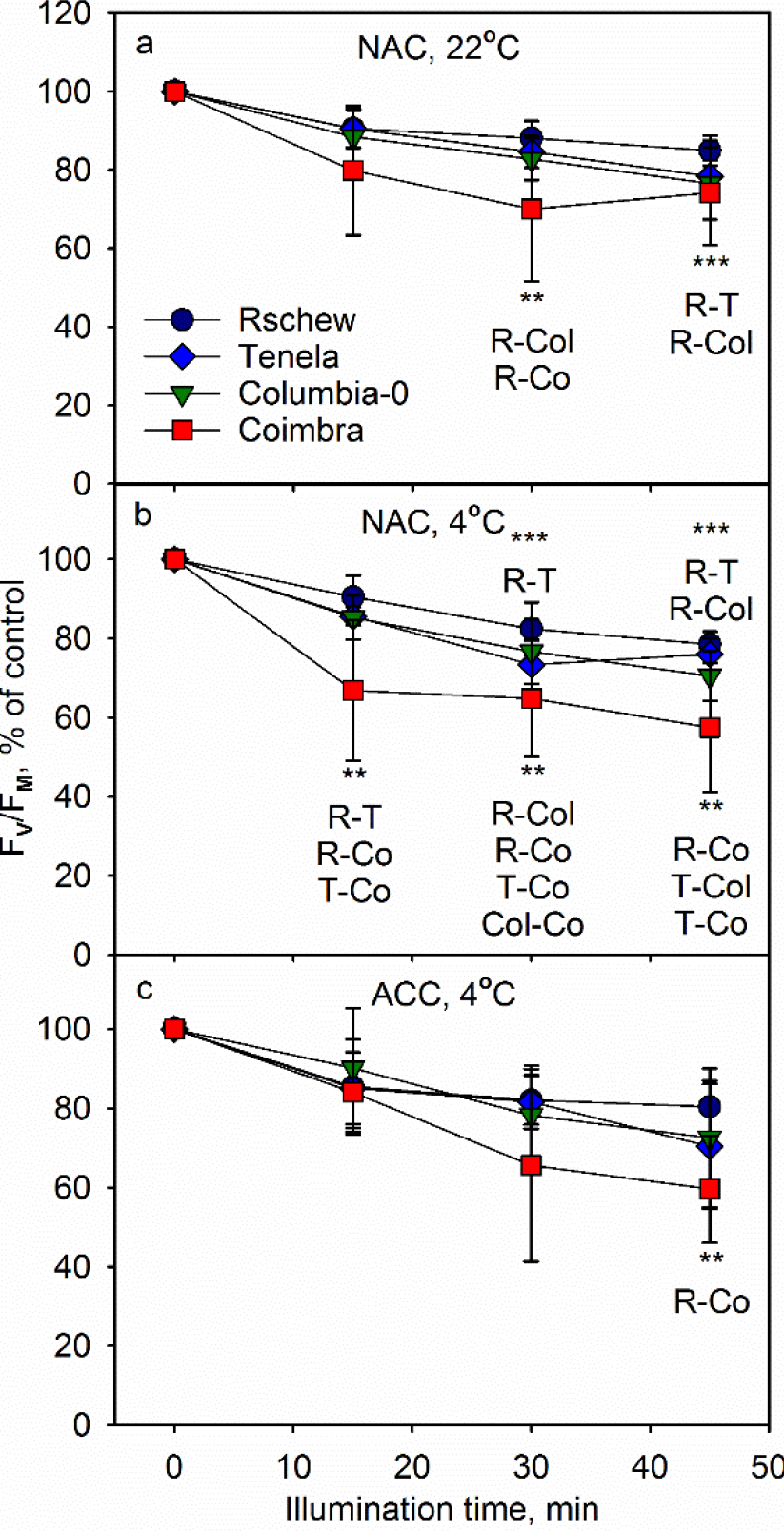
Photoinhibition, quantified by the chlorophyll *a* fluorescence parameter F_V_/F_M_, at 22 °C (a) or at 4 °C (b–c) in the absence of lincomycin. F_V_/F_M_ was measured from detached leaves of four non-acclimated (NAC; a–b) or cold-acclimated (ACC; c) *A. thaliana* accessions, at different time points during the 45-min illumination (PPFD 2000 μmol m^−2^s^−1^) and subsequent 30-min dark incubation. The error bars show standard deviations (SD) from at least three biological replicates. Statistically significant differences at any time-point between the indicated accessions are marked with *** (*P* < 0.01) or ** (*P* < 0.05). The control values of F_V_/F_M_ (±SD) were 0.82 (0.03), 0.80 (0.03), 0.82 (0.03) and 0.76 (0.05) for Rschew (R), Tenela (T), Columbia-0 (Col) and Coimbra (Co), respectively, in (a), 0.82 (0.03), 0.82 (0.03), 0.81 (0.04) and 0.79 (0.06) in (b), and 0.82 (0.03), 0.79 (0.04), 0.77 (0.08) and 0.79 (0.06) in (c)

### 3.2. Assaying photoinhibition of PSII, in the absence of repair, by chlorophyll *a* fluorescence

To see if the observed differences in photoinhibition were due to differences in the rate of damage to PSII or in the rate of repair, we repeated the experiments in the presence of lincomycin. In addition, the absence of the repair allows a decline in PSII activity (quantified by F_V_/F_M_) to be fitted to the first order reaction equation for calculation of the rate constant of photoinhibition of PSII (k_PI_).

Similarly as observed without lincomycin (Fig. 1), we found that Coimbra and Rschew were, respectively, the least and the most tolerant to photoinhibition, both at 22 °C and at 4 °C (Fig. 2A–B, see Table 1 for the k_PI_ values). However, in contrast to the experiments in which the repair was allowed to function, F_V_/F_M_ values declined more rapidly at 22 °C than at 4 °C in Columbia-0 (*P* = 0.02) and Coimbra (*P* = 0.05) (Fig. 2A–B). In Rschew and Tenela the differences were not statistically significant. In all four accessions, photoinhibition at 4 °C proceeded more slowly in ACC plants than in NAC plants (Fig. 2B–C) Accordingly, the k_PI_ values of ACC Rschew, Tenela and Coimbra were circa 77 % and of ACC Columbia-0 84 % of those of NAC plants (Table 1). The difference was significant in the case of Rschew (*P* = 0.0002) and Tenela (*P* = 0.01). The differences in photoinhibition tolerance were no longer statistically significant among the ACC accessions (Fig. 2C, Table 1).

**Fig. 2.**
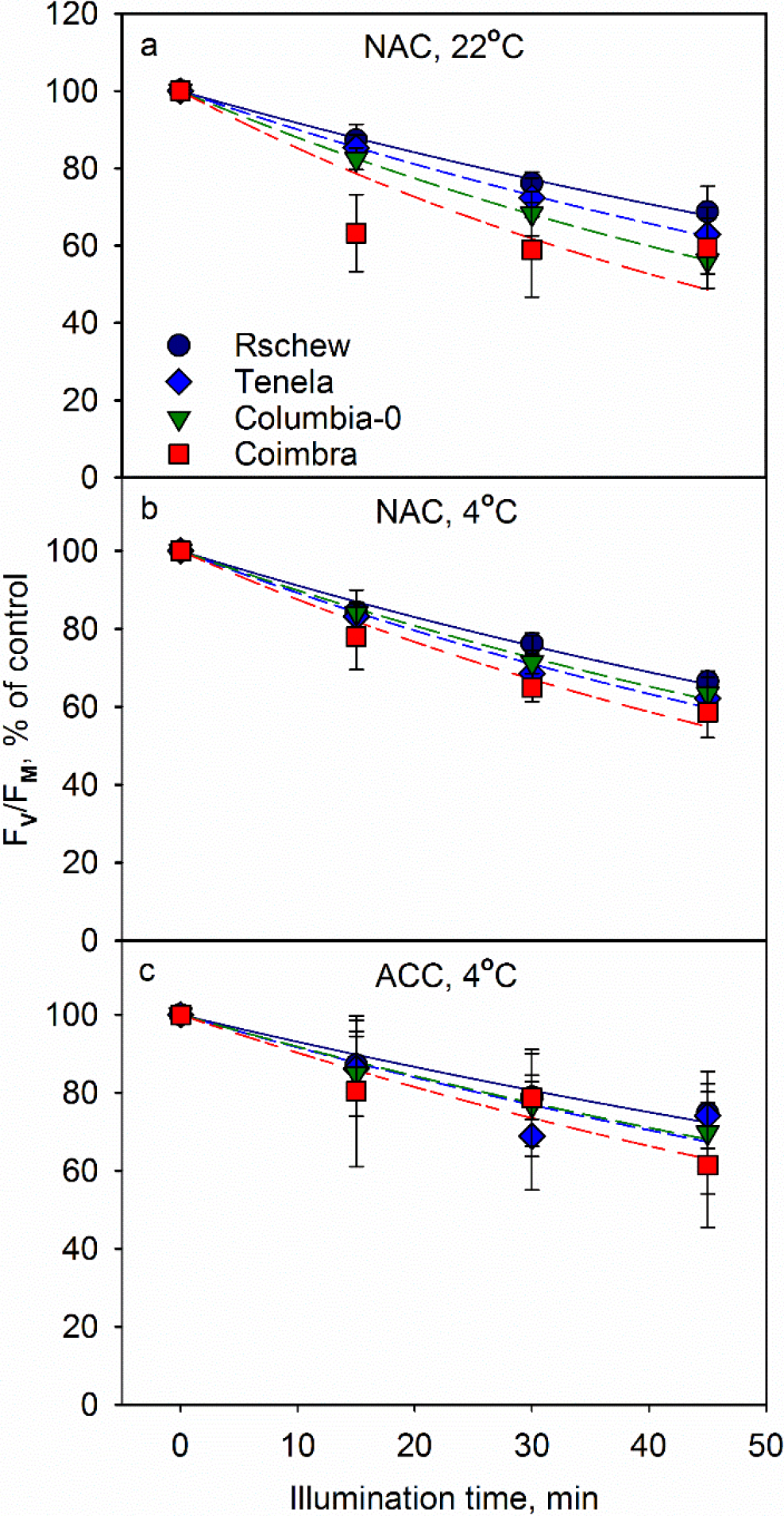
Photoinhibition, quantified by the chlorophyll *a* fluorescence parameter F_V_/F_M_, at 22 °C (a) or at 4 °C (b–c) in the presence of lincomycin. F_V_/F_M_ was measured from detached leaves of four non-acclimated (NAC; a–b) or cold-acclimated (ACC; c) *A. thaliana* accessions, at different time points during the 45-min illumination (PPFD 2000 μmol m^−2^s^−1^) and subsequent 30-min dark incubation. The error bars show SD from at least three biological replicates. The control values of F_V_/F_M_ (±SD) were 0.82 (0.03), 0.79 (0.05), 0.83 (0.01) and 0.78 (0.04) for Rschew, Tenela, Columbia-0 and Coimbra, respectively, in (a), 0.80 (0.01), 0.77 (0.02), 0.81 (0.03) and 0.79 (0.03) in (b) and 0.82 (0.03), 0.77 (0.07), 0.79 (0.06) and 0.79 (0.06) in (c)

**Table 1.** The rate constant of photoinhibition (k_PI_) values in min^−1^ calculated by fitting the decline in F_V_/F_M_ (Fig. 2) or in PSII-dependent oxygen evolution (Fig. 3) during the 45-min illumination at 22 °C or at 4 °C, to the first order reaction equation in NAC and ACC *A. thaliana* accessions. SD values (in parentheses) were calculated from at least three biological replicates. Statistically significant differences (*P* < 0.05) between accessions are indicated with lower-case letters and between treatment groups (between NAC 22 °C and NAC 4 °C, or between NAC 4 °C and ACC 4 °C) with upper-case letters. Significances of the differences between different accessions are shown only within the same treatment group. Significances of the differences between fluorescence and oxygen evolution data were not calculated

| | $k_{PI}$, $\text{min}^{-1}$ (SD) | | | | | |
| --- | --- | --- | --- | --- | --- | --- |
|  | Non-acclimated, at 22°C |  | Non-acclimated, at 4°C |  | Cold-acclimated, at 4°C |  |
|  | Fluorescence | Oxygen evolution | Fluorescence | Oxygen evolution | Fluorescence | Oxygen evolution |
| **Rschew** | 0.0086<sup>a,A</sup><br>(0.0011) | 0.0194<sup>a,A</sup><br>(0.0044) | 0.0093<sup>a,A</sup><br>(0.0008) | 0.0124<sup>a,B</sup><br>(0.0044) | 0.0072<sup>a,B</sup><br>(0.0016) | 0.0155<sup>ab,B</sup><br>(0.0048) |
| **Tenela** | 0.0105<sup>b,A</sup><br>(0.0012) | 0.0210<sup>ab,A</sup><br>(0.0024) | 0.0115<sup>bc,A</sup><br>(0.0013) | 0.0192<sup>b,A</sup><br>(0.0047) | 0.0088<sup>a,B</sup><br>(0.0034) | 0.0220<sup>a,A</sup><br>(0.0048) |
| **Columbia-0** | 0.0128<sup>c,A</sup><br>(0.0014) | 0.0247<sup>b,A</sup><br>(0.0036) | 0.0111<sup>b,B</sup><br>(0.0027) | 0.0179<sup>ab,B</sup><br>(0.0059) | 0.0093<sup>a,B</sup><br>(0.0042) | 0.0128<sup>b,B</sup><br>(0.0044) |
| **Coimbra** | 0.0160<sup>d,A</sup><br>(0.0011) | 0.0241<sup>ab,A</sup><br>(0.0051) | 0.0135<sup>c,B</sup><br>(0.0027) | 0.0226<sup>b,A</sup><br>(0.0080) | 0.0105<sup>a,B</sup><br>(0.0053) | 0.0124<sup>b,B</sup><br>(0.0044) |

**Fig. 3.**
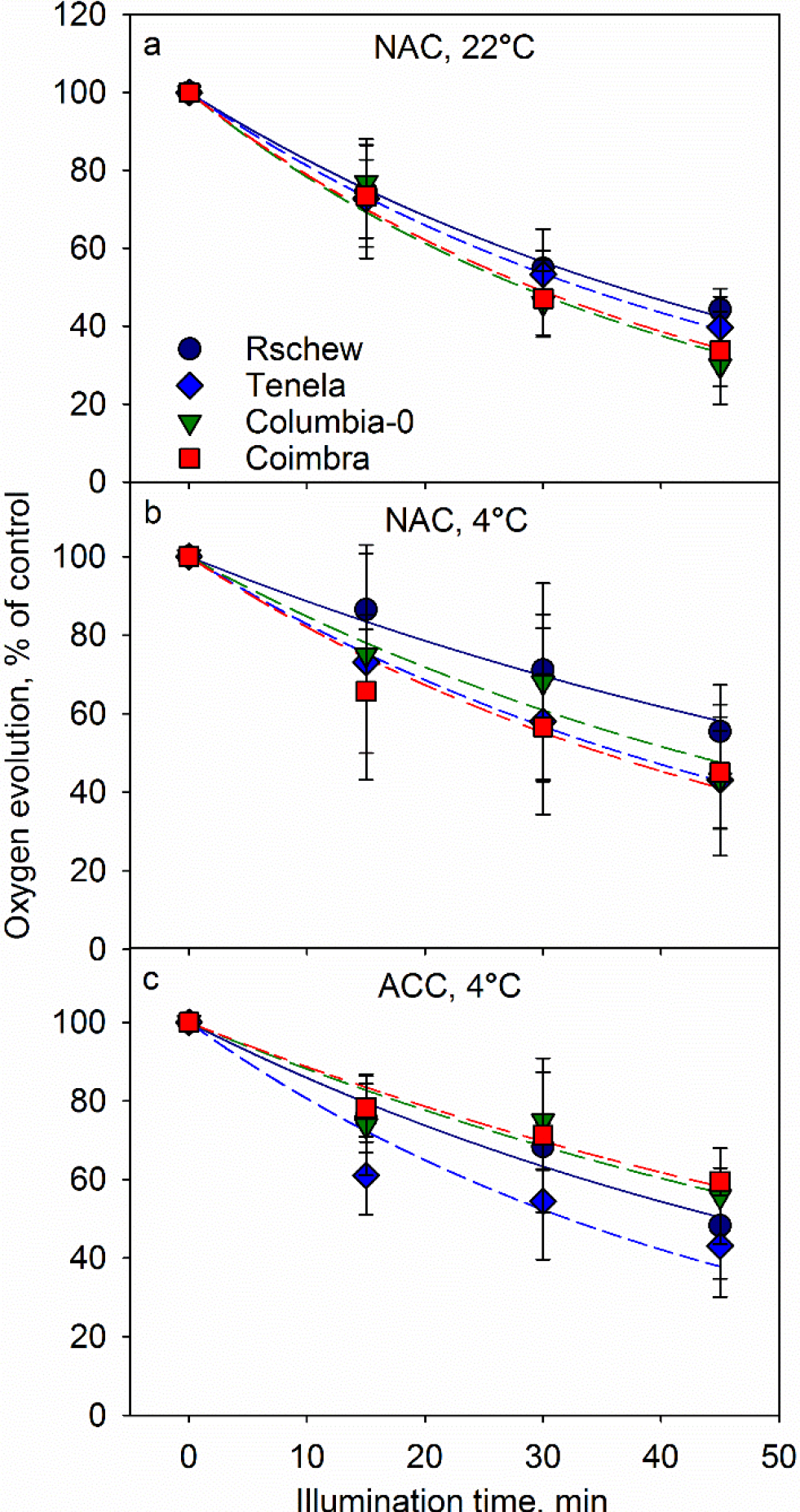
Photoinhibition, quantified by the light-saturated oxygen evolution of PSII in the presence of an artificial electron acceptor, at 22 °C (a) or at 4 °C (b–c) in the presence of lincomycin. PSII activity was measured from thylakoid membranes isolated from leaves of four non-acclimated (NAC; a–b) or cold-acclimated (ACC; c) *A. thaliana* accessions, at different time points during the 45-min illumination (PPFD 2000 μmol m^−2^s^−1^). The error bars show SD from at least four biological replicates. The control values in μmol O_2_ (mg chl)^−1^ h^−1^ (±SD) were 160 (37.5), 162 (26.4), 180 (20.2) and 132 (31.2) for Rschew, Tenela, Columbia-0 and Coimbra, respectively, in (a), 169 (67.6), 148 (67.1), 162 (37.4) and 135 (24.7) in (b) and 210 (21.7), 186 (22.9), 164 (28.7) and 162 (37) in (c)

### 3.3. Assaying photoinhibition of PSII, in the absence of repair, by oxygen evolution

To test whether the results are universal or specific to a particular method of quantification of photoinhibition, we next assayed photoinhibition of PSII by measuring the light-saturated rate of oxygen evolution in the presence of an artificial electron acceptor from thylakoid membranes isolated from the illuminated leaves.

Photoinhibition appeared faster when measured with oxygen evolution compared to that measured with the fluorescence parameter F_V_/F_M_ (c.f. Figs. 2–3; Table 1). However, similarly to the fluorescence data, we found that Rschew was the most tolerant accession and Coimbra the least, at both 4 °C and 22 °C (Table 1). Furthermore, photoinhibition proceeded more slowly at 4 °C than at 22 °C in all the accessions (Fig. 3A–B), although the differences in the k_PI_ values (Table 1) were significant (*P* < 0.05) only for Rschew and Columbia-0. Cold-acclimation alleviated photoinhibition at 4 °C statistically significantly only in Coimbra (Fig. 3B–C). In Rschew and Tenela, the differences between the k_PI_ values for ACC and NAC plants were not statistically significant.

### 3.4. Physiological parameters of cold-acclimated and non-acclimated plants

To find the factors affecting the different photoinhibition tolerances of the accessions, we measured chlorophyll content, epidermal flavonols, photochemical reflectance index (PRI) and the rate of CO_2_ assimilation of intact leaves of NAC and ACC accessions, by using non-invasive instruments. Chlorophyll contents, measured per leaf area with an optical method, were only little affected by temperature, age or accession. Cold-acclimation caused slight decrease of the chlorophyll content after seven days at 4 °C only in Columbia-0 (Fig. 4A), and even in this accession, no statistically significant difference was observed after 14 days of the cold-acclimation. In NAC Rschew the chlorophyll content slightly increased during the 14 day time-frame of the experiment (Fig. 4A). The relative differences in the amounts of chlorophyll between the accessions (Rschew > Columbia-0 > Tenela > Coimbra) stayed the same in all the treatment groups (Fig. 4A).

**Fig. 4.**
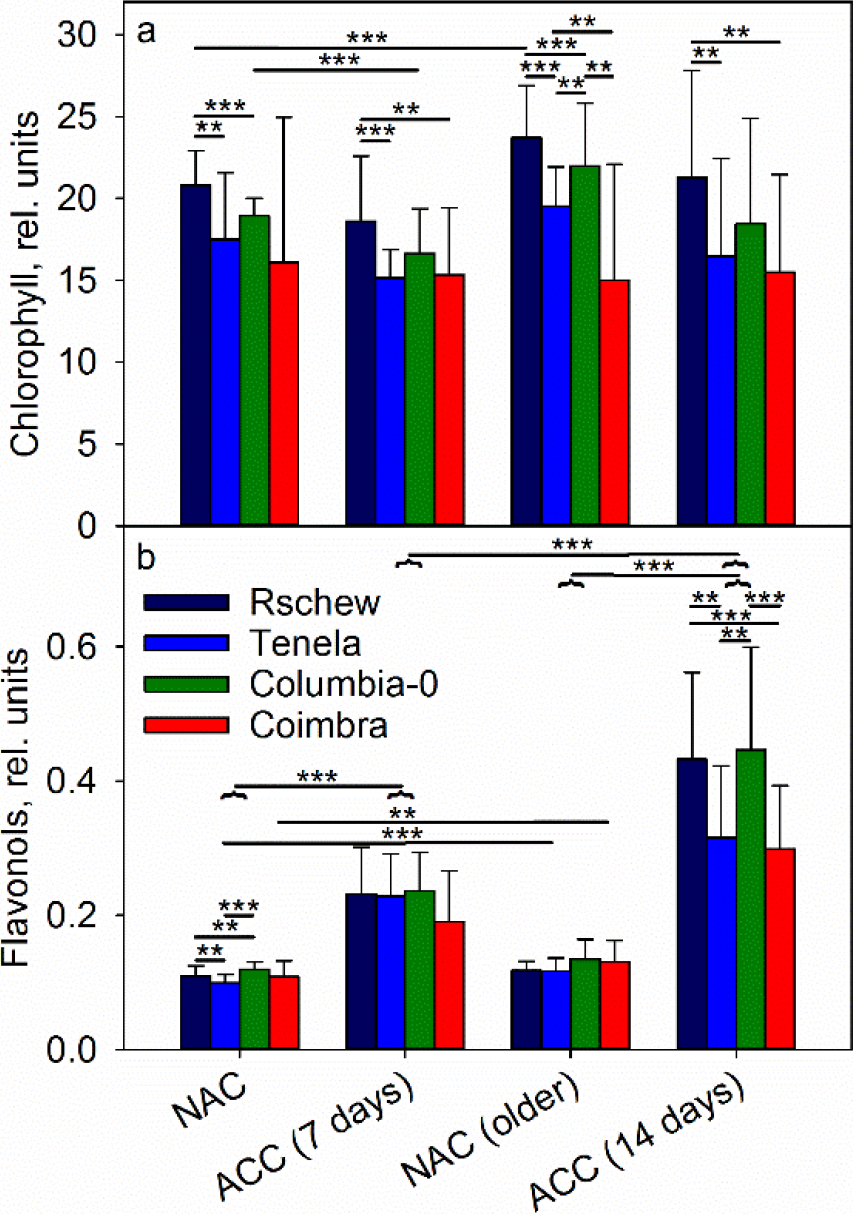
Chlorophyll (a) and flavonol (b) contents measured from intact leaves of *A. thaliana* accessions (Rschew, Tenela, Columbia-0 and Coimbra) after 7 or 14 days, as indicated, of cold-acclimation at 4 °C (ACC) or from non-acclimated (NAC) control plants of similar ages. The error bars show SD values from at least four biological replicates. Statistically significant differences are marked with ** (*P* < 0.05) or *** (*P* < 0.01) on top of the horizontal lines that show between which samples the difference is significant; the horizontal curly bracket indicates a whole group of four accessions. The significance of the differences between different accessions are shown only within the same treatment group, and significances between NAC and ACC plants are shown only between the corresponding age groups

Contrary to chlorophylls, epidermal flavonols increased significantly during the cold-acclimation, both after 7 and 14 days, in all the four accessions (Fig. 4B). In Tenela and Coimbra the cold-acclimation-induced increase in epidermal flavonols was lower than in Rschew and Columbia-0, and the difference was more obvious after 14 days of cold-acclimation (Fig. 4B). In Tenela and Coimbra, flavonols also increased when the NAC plants grew older (Fig. 4B).

Even though PSII activity declined by about 20 % during the 45-min high light illumination in the absence of lincomycin at 22 °C (Fig. 1A), a similar illumination treatment did not cause a decline in the net rate of CO_2_ assimilation, measured per leaf area (Fig. 5). The maximum assimilation rates varied between the accessions, and the values (Fig. 5) and the chlorophyll content of the leaves (Fig. 4A) showed similar order between the accessions, with Rschew highest and Coimbra and Columbia-0 low, especially in NAC plants. Cold-acclimation caused a decrease in the rate of CO_2_ assimilation in high light at 22 °C in all four accessions (Fig. 5).

**Fig. 5.**
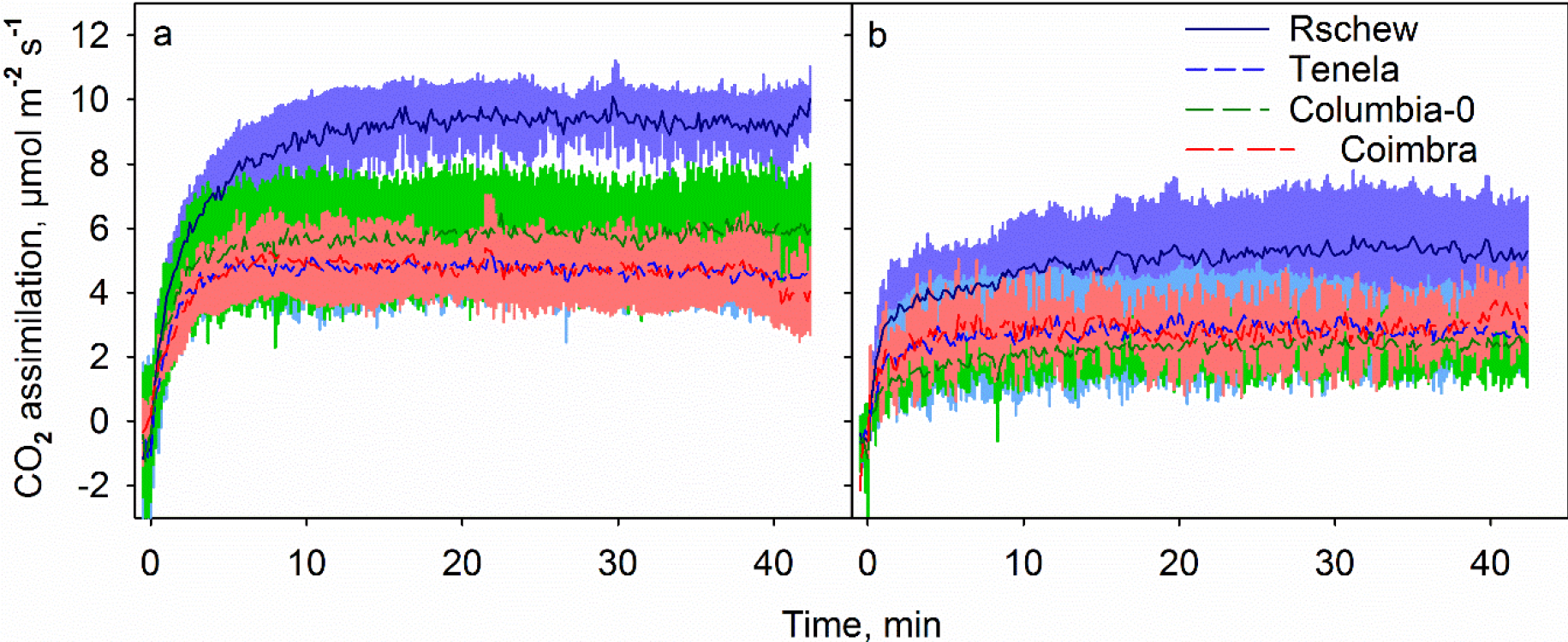
Net CO_2_ assimilation rates at 22 °C during 42-min illumination in high light (PPFD 2000 μmol m^−2^s^−1^), measured from intact light-acclimated leaves of non-acclimated (a) or cold-acclimated (b) *A. thaliana* accessions (Rschew, Tenela, Columbia-0 and Coimbra). The lines show averages and the colored areas show SD values from at least three biological replicates

### 3.5. ^1^O_2_ production in thylakoid membranes

To see if the physiological differences of the accessions lead to differences in the production of ^1^O_2_, we measured the rate of ^1^O_2_ production in isolated thylakoid membranes of NAC and ACC plants at 20 °C and at 4 °C. At 20 °C, thylakoids isolated from ACC plants produced more ^1^O_2_ than those isolated from NAC plants but the difference was statistically significant only in the case of Coimbra (Fig. 6). In both NAC and ACC thylakoids, production of ^1^O_2_ was slightly slower at 4 °C than at 20 °C, but statistically significant difference in ^1^O_2_ production was observed only in ACC Coimbra. Interestingly, the production of ^1^O_2_ by ACC thylakoids at 4 °C did not significantly differ from that of NAC thylakoids (Fig. 6).

**Fig. 6.**
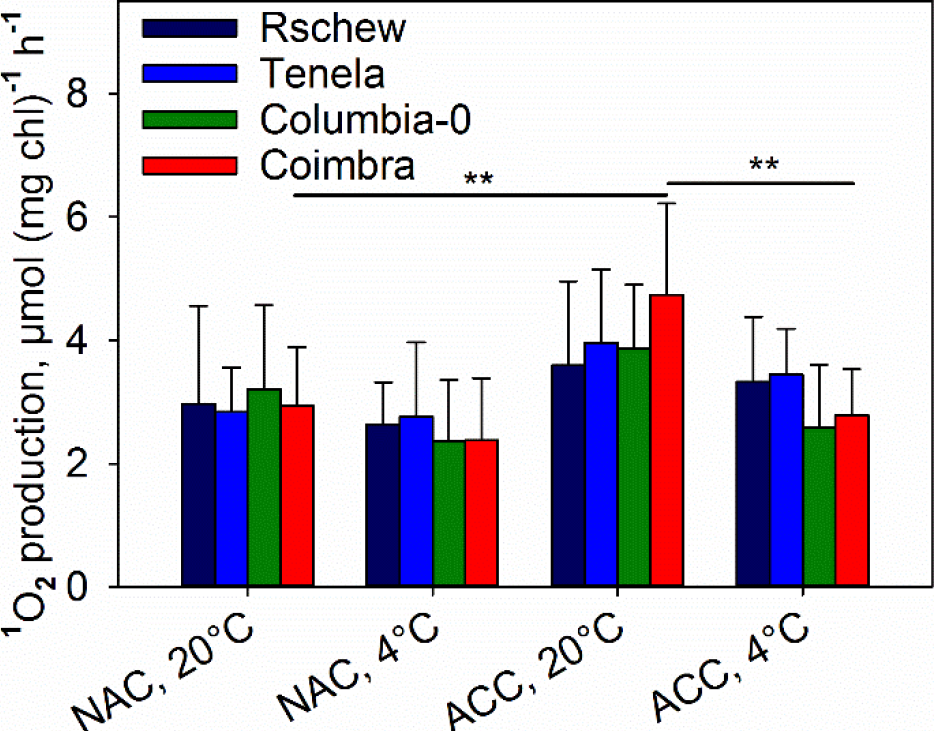
Production of ^1^O_2_ in high light (PPFD 4000 μmol m^−2^s^−1^) at 20 °C or 4 °C in thylakoid membranes isolated from non-acclimated (NAC) or cold-acclimated (ACC) *A. thaliana* accessions (Rschew, Tenela, Columbia-0 and Coimbra). ^1^O_2_ was measured with a histidine-based method. The error bars show SD values from at least four biological replicates. Statistically significant differences are marked with ** (*P* < 0.05). The significances of the differences between different accessions are shown only within the same treatment group, and significances between NAC and ACC plants are shown only between the corresponding temperatures

Modulation of charge recombination reactions in PSII by cold-acclimation has been suggested to lead to diminished ^1^O_2_ production (Ivanov 2008). To assess the recombination reactions, thermoluminescence was measured from the same thylakoid membranes used for the ^1^O_2_ assay. In all samples, the peak of the B-band, measured in the absence of DCMU, was between 25 °C and 28 °C whereas the Q-band (measured in the presence of DCMU) peaked around 7–9 °C with no systematic differences between accessions or between NAC and ACC plants (Fig. 7). However, the thermoluminescence yield of the B-band was 4–46 % and the yield of the Q-band 26–46 % lower for the ACC than NAC plants (relative peak intensities are listed in the legend of Fig. 7).

**Fig. 7.**
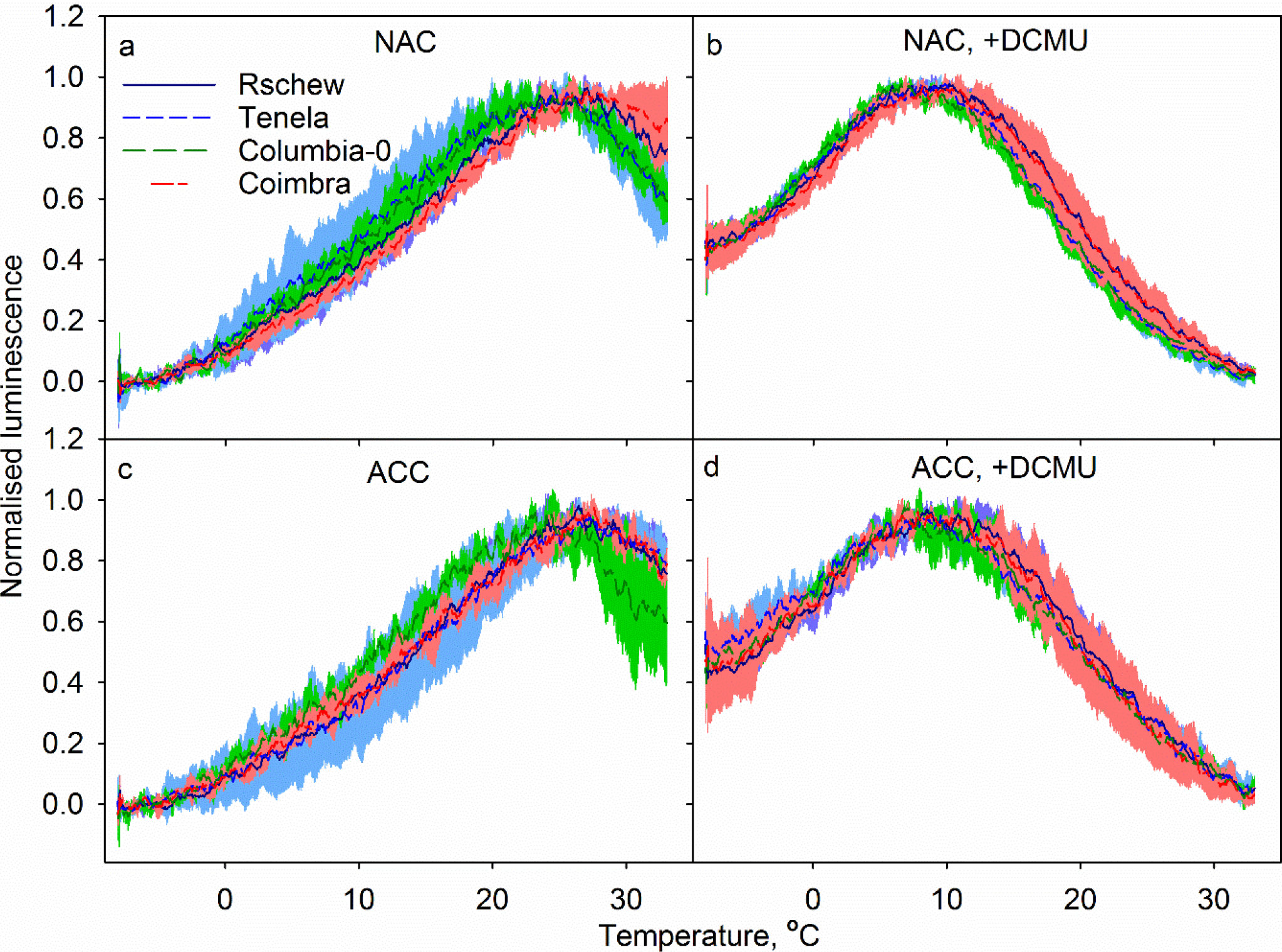
Normalized thermoluminescence bands in the absence (a, c) or in the presence of DCMU (b, d) measured from thylakoid membranes of non-acclimated (NAC; a–b) or cold-acclimated (ACC; c–d) *A. thaliana* accessions (Rschew, Tenela, Columbia-0 and Coimbra). The colored areas show SD values from at least three biological replicates. Maximum luminescence intensities (arbitrary units; ±SD) were 2.1 (0.60), 1.8 (0.45), 1.7 (0.15) and 2.8 (0.21) for Rschew, Tenela, Columbia-0 and Coimbra, respectively, in (a), 2.5 (0.77), 2.2 (0.23), 2.2 (0.17) and 2.6 (0.39) in (b), 2.0 (0.26), 1.5 (0.20), 1.4 (0.25) and 1.5 (0.2) in (c) and 1.8 (0.65), 1.2 (0.41), 1.6 (0.16) and 1.9 (0.08) in (d)

## 4. Discussion

### 4.1. The damaging reaction of photoinhibition *in vivo* is slower at 4 °C than at 22 °C

Temperature affects the rate of the damaging reaction of photoinhibition. To get a comprehensive understanding, we used two parallel methods for measuring photoinhibition of PSII, i.e. the chlorophyll *a* fluorescence parameter, F_V_/F_M_, and PSII-dependent oxygen evolution measured from thylakoids isolated from the treated leaves. Comparison of the present data obtained from the NAC plants of all accessions show a clear trend of slower photoinhibition at 4 °C than at 22 °C (Table 1), although the fluorescence data do not show statistically significant differences in accessions Rschew and Tenela (Table 1; Fig. 2). A slight increase in the rate of the damaging reaction at higher temperatures corroborates earlier *in vitro* (Aro et al. 1990; Tyystjärvi et al. 1994) and *in vivo* (Tyystjärvi 2013) results. These findings suggest that the excitation pressure hypothesis (see Kornyeyev et al., 2003) is not a valid explanation for photoinhibition of PSII.

The data discussed above contrast with the findings of Tsonev and Hikosaka (2003) and Kornyeyev et al. (2003) who observed faster decline in F_V_/F_M_ at low than at optimal temperature. It is possible that the discrepancy originates from the methods used for assessing photoinhibition. F_V_/F_M_ may reflect low-temperature-induced fluorescence quenching that does not relax during a typical dark-adaptation time (20–60 min). Such slowly relaxing F_V_/F_M_ might be related to the sustained NPQ that develops in some evergreen species at low temperatures (for a review, see Verhoeven 2014) and might be important also in *A. thaliana* (Malnoë et al. 2018). Therefore, especially in studies of cold-acclimation, use of oxygen evolution as a proxy for photoinhibition is advisable.

Due to the fact that light absorption continues but carbon fixation and other enzymatic reactions slow down, light can possess a severe stress for plants (see e.g. Alboresi et al. 2011), even though the rate of light-induced damage to PSII is not accelerated at low temperatures (Table 1; Fig. 3). As low temperature commonly slows down the enzymatic reactions of concurrent recovery of photoinhibitory damage (Greer et al. 1986; Aro et al. 1990) and ROS produced at low temperature may specifically inhibit translation in the chloroplast (Kojima et al. 2009), net loss of PSII activity during illumination at temperatures below the optimum occurs (Tyystjärvi and Aro 1990; Öquist et al. 1993; Allakhverdiev and Murata 2004). The results of the present study confirm faster loss of PSII activity at 4 °C than at 22 °C when no protein synthesis inhibitor was used (Fig. 1A–B) that corresponds, taking into account the rate of the damage (Table 1), to slower repair cycle of PSII at 4 °C than at 22 °C.

### 4.2. Cold-acclimation may increase the tolerance against photoinhibition in *A. thaliana* accessions

Plants’ capacity to cold-acclimate vary greatly (e.g. Hannah et al. 2006, Hasdai et al. 2006, Oakley et al. 2018) and a correlation exists between the distance of a species or accession from the equator and their capacity to gain cold-tolerance (Hannah et al. 2006; Mishra et al. 2011; Lukas et al. 2014). The LT_50_ (lethal temperatures at which 50 % of tissue damage occurs as measured by electrolyte leakage) has been reported to be −5.7 °C, −7.7 °C, −5.2 °C and 4.6 °C for Rschew, Tenela, Columbia-0 and Coimbra, respectively (Hannah et al. 2006; Mishra et al. 2011). Two weeks of cold-acclimation at 4 °C can significantly increase the cold-tolerance in Rschew, Tenela and Columbia-0 (to −10.6 °C, −12.2 °C and −10.3 °C, respectively), however, Coimbra is able to decrease the LT50 only by about 1.5 °C (Hannah et al. 2006).

Cold-acclimation slowed down photoinhibition of PSII by slowing down the damaging reaction of photoinhibition of PSII in all accessions when measured with F_V_/F_M_, and the difference was statistically significant in the two most cold-tolerant accessions Rschew and Tenela (Fig. 2; Table 1), similarly as reported previously for cyanobacteria (Vonshak and Novoplansky 2008). Oxygen evolution data, in turn, showed protection by cold-acclimation only in the accessions Columbia-0 and Coimbra, which can be considered as intermediate cold-tolerant and cold-susceptible, respectively.

The results suggest that cold-acclimation can harness plants against photoinhibition. However, in the accessions used in the present study the differences were not big, and Rschew and Tenela were actually more susceptible to photoinhibition after cold-acclimation, when oxygen evolution was used to quantify photoinhibition, though the difference was not statistically significant (Fig. 3; Table 1). This may indicate that the acclimation that induces cold-tolerance in *A. thaliana* does not primarily provide protection against PSII damage. It should be noted, however, that in the present study plants were grown under moderate light (PPFD 100 μmol m^−2^ s^−1^), which may impair their capacity to respond to high-light-treatments.

In addition, even though cold-acclimation seemed to attenuate photoinhibition also in the absence of lincomycin, the differences were neither large nor statistically significant (Fig. 1B–C), suggesting that in the natural accessions used, enhancement of the repair reactions of PSII is not a prominent factor in cold-tolerance. In fact, photoinhibition (or decrease in photosynthesis in more general) has been suggested not to be the cause of decreased productivity but regulation responding to diminished sink demand (see Adams et al. 2013). Accordingly, CO_2_ assimilation at 22 °C (Fig. 5) did not correlate with the amount of photoinhibition in any NAC accessions (Figs. 1–3). In addition, net CO_2_ assimilation was slower in cold-acclimated leaves than in their NAC counterparts; however, carbon fixation at 4 °C was not measured. The lack of change in the rate of carbon fixation under strong light also suggests that *A. thaliana* leaves contain more PSII than needed to saturate the needs of the carbon fixation reactions in high light (see also Chow 2001).

### 4.3. Decrease in thermoluminescence yield did not lower ^1^O_2_ production

What causes the cold-acclimation-induced alleviation of photoinhibition? ^1^O_2_ is produced mainly in the recombination reactions of PSII, and it has been suggested to cause the photodamage to PSII (e.g. Vass 2012). Increased production of ^1^O_2_ in the *npq1lut2* mutant of *A. thaliana* leads to oxidative damage to thylakoid proteins in high light at 10 °C (Alboresi et al. 2011). In different plants species including *A. thaliana* (Janda et al. 2000; Ivanov et al. 2001; Sane et al. 2003), cold-acclimation causes a decrease in the redox gap between the Q_A_ and Q_B_ electron acceptors of PSII, which favors an indirect, non-radiative recombination route (Rappaport and Lavergne 2009) for the elimination of the charge in Q_A_^−^(Ivanov 2008). The indirect route does not produce ^1^O_2_ and, therefore, the observed decrease in photoinhibition caused by cold-acclimation might be due to a decrease in ^1^O_2_ production (for a more comprehensive discussion about the relationship between recombination reactions and ^1^O_2_, see Krieger-Liszkay et al. 2008; Vass and Cser 2009). Accordingly, Ramel et al. (2012) observed a decrease in both production of ^1^O_2_ and oxidation of beta-carotene after 99 h at 7 °C. However, ^1^O_2_ was measured with Singlet Oxygen Sensor Green in white light, a condition that has been shown to induce ^1^O_2_ production by the sensor itself (e.g. Ragás et al. 2009).

We did not observe shifts in the peak temperatures of the thermoluminescence bands (Fig. 7). Discrepancy with the earlier data (e.g. Sane et al. 2003) may result from different durations of the cold-acclimation treatments. We cold-acclimated the *A. thaliana* accessions for 14 days which is enough for cold-tolerance (e.g. Mishra et al. 2011; Mishra et al. 2014), but stronger alleviation of photoinhibition was reported when each leaf developed at low temperature (Strand et al. 1999; Gray et al. 2003). However, we did observe a decrease in the thermoluminescence yield for both B- and Q-bands, resembling that reported by Ivanov et al. (2001) and Sane et al. (2003). The intensity of the B-band decreased only little, ~4 %, in Rschew but the decrease was more remarkably in Tenela (18 %), Columbia-0 (20 %) and Coimbra (46 %). We also observed strong alleviation of photoinhibition in the cold-acclimated Coimbra. However, in other accessions or in the Q-band, the decrease in the intensity did not correlate with the rate of photoinhibition (Fig. 7, Table 1). Furthermore, cold-acclimation did not cause a decrease in the capacity of the thylakoids to produce ^1^O_2_ during illumination (Fig. 6).

If the rate of repair is insignificant at low temperatures, then the results from illumination treatments without an inhibitor of (chloroplast) protein synthesis could be taken to represent the amount of damage to PSII. However, in the present data, addition of lincomycin clearly increased photoinhibition, indicating that the repair cycle is active even at 4 °C in NAC plants (Figs. 1–2). Also the finding that genes coding for a protease involved in PSII repair cycle (FtsH) are up-regulated at low temperature (Soitamo et al. 2008) support this view. Consequently, it remains unclear whether the (proposed) diminished ^1^O_2_ production by modified recombination reactions (Ivanov at al. 2001; Sane et al. 2003) decreases PSII damage or protects the repair reactions (Kojima et al. 2009; Nishiyama et al. 2011).

### 4.4. Sensitivity to ROS does not equal sensitivity to photoinhibition of PSII

The roles of ROS in the cold-acclimation-induced protection against photoinhibition are interesting also with regard to the mechanism of photoinhibition. The k_PI_ values of the four accessions, measured by both F_V_/F_M_ and oxygen evolution, were mostly in the order Rschew < Tenela < Columbia-0 < Coimbra (Table 1). Furthermore, Coimbra was clearly more susceptible to the damage caused by high light than the other accessions when the PSII repair cycle was allowed to run (Fig. 1). Thus, the two cold-tolerant accessions were somewhat more tolerant to photoinhibition than the other two accessions (Columbia-0 and Coimbra). Previously, it has been shown that Tenela is sensitive to oxidative stress (Brosché et al. 2010), which is probably linked to increased production of hydrogen peroxide at low temperatures (Distelbarth et al. 2013). However, in our study non-acclimated Tenela was not particularly sensitive to photoinhibition of PSII (Figs. 1–3), which supports the suggestion that oxidative stress and photoinhibition may not be related (Hakkila et al. 2013). Also, the rate of light-induced damage to PSII may not be defined by ROS but rather direct light absorption by the oxygen evolving complex of PSII (Hakala et al. 2005; Ohnishi et al. 2005). Furthermore, the production of ^1^O_2_ by isolated thylakoid membranes did not correlate with the k_PI_ values (Fig. 8A), suggesting that ^1^O_2_ is not a decisive factor in photoinhibition tolerance of *A. thaliana* accessions.

**Fig. 8.**
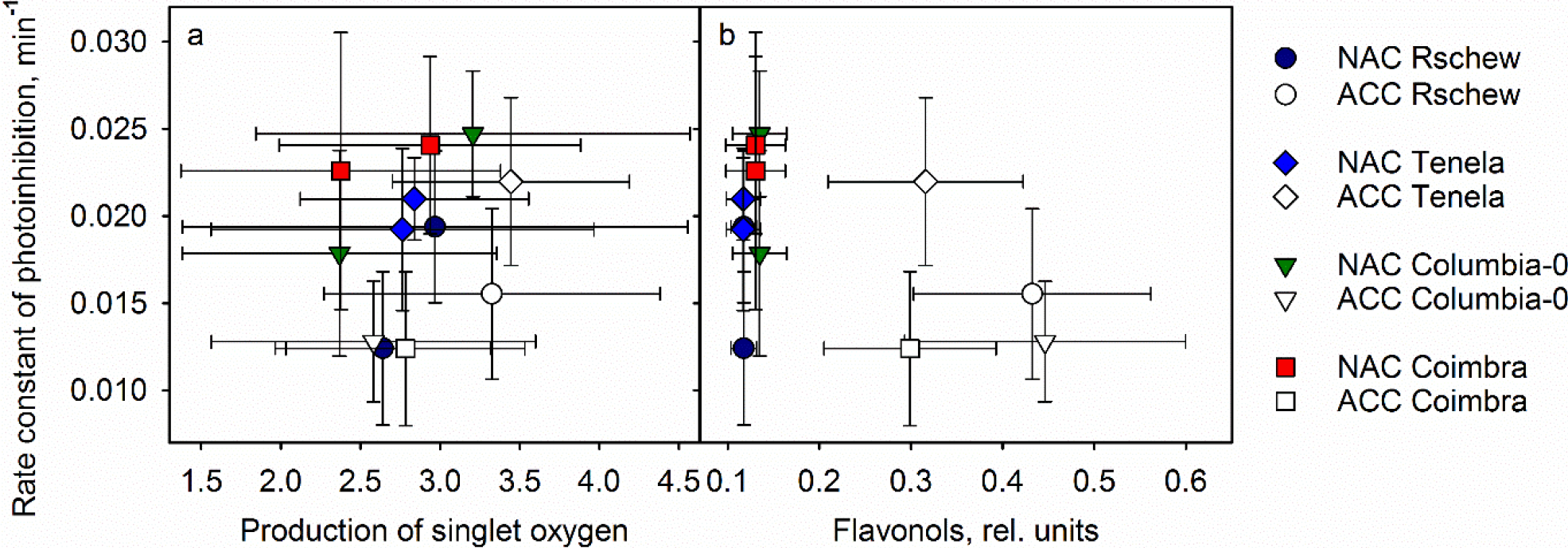
Rate constant of photoinhibition (quantified as a loss of oxygen evolution capacity of PSII) plotted against production of ^1^O_2_ by thylakoid membranes (a) and the amount of epidermal flavonols (b) in four *A. thaliana* accessions (Rschew, Tenela, Columbia-0 and Coimbra) after 14 days of cold acclimation at 4 °C (ACC) or from control plants of similar ages grown at 21 °C (NAC). The data are from Figs. 4 and 6 and from Table 1

Rather than reflecting the ability to resist direct damage by ROS, the sensitivity to oxidative stress may also be governed by a genetic program (Brosché et al. 2010). Hydrogen peroxide, which has a relatively long lifetime in cells, is an important signaling molecule (for a review see e.g. Černý et al. 2018). Further complications in the roles of ROS in cold-acclimation are exemplified by discrepant findings about the relationship between ROS metabolism and freezing tolerance (Distelbarth et al. 2013; Hashempour et al. 2014).

### 4.5. Do flavonols quench ^1^O_2_ and/or protect plants from photoinhibition?

Synthesis of many flavonol species is induced in coldness (e.g. Schulz et al. 2015) as well as in UV light (e.g. Hectors et al. 2014). Flavonols are able to physically and chemically quench ^1^O_2_ *in vitro* (Tournaire et al. 1993), and it has been suggested that they function as ^1^O_2_ quenchers also *in vivo* (Pollastri and Tattini 2011; Majer et al. 2014). Accordingly, Havaux and Kloppstech (2001) reported that mutants unable to synthetize anthocyanins and flavonols had lower F_V_/F_M_ values and more lipid peroxidation at low temperatures than wild-type plants.

Although in Columbia-0 and Coimbra one can see a positive correlation between flavonols and the rate constant of photoinhibition, suggesting that quenching of ^1^O_2_ by flavonols can in some accessions protect against photoinhibition, general correlation was not found (Fig. 8B). In addition, even though cold-acclimation increased the amount of flavonols in all accessions tested here (Fig. 4), the rate of ^1^O_2_ production by isolated thylakoid membranes did not depend on the flavonol content of the leaf (Fig. 6).

Of course, the optical method used in the present study mainly reflects the amount of epidermal flavonols. Therefore, the role of flavonols, both inside and outside of chloroplasts, in cold-tolerance requires more research. Flavonols may be important as freezing inhibitors (Isshiki et al. 2014; Korn et al. 2008). Flavonols may also have regulatory roles (Yin et al. 2014).

## Acknowledgements

This work was supported by Academy of Finland (grant 307335; to E.T.), Turku University Foundation (12353; to H.M.), University of Turku Graduate School (to H.M.), Vilho, Yrjö and Kalle Väisälä Foundation (to H.M.), European Union COST Action (TD1102; to E.T., H.M. and K.B.M.), the Ministry of Education, Youth and Sports of CR within the National Sustainability Program I (NPU I, LO1415; to K.B.M., A.M., K.A. and D.S.) and the project “SustES − Adaptation strategies for sustainable ecosystem services and food security under adverse environmental conditions” (CZ.02.1.01/0.0/0.0/16_019/0000797; to K.B.M.).

